# The effects of alternative rabbit control methods on feral cat activity in an open, semi-arid landscape

**DOI:** 10.1101/2023.12.03.569789

**Authors:** Jeroen Jansen, Sebastien Comte, Abbey T Dean, Geoff Axford, Katherine E Moseby, David E Peacock, Robert Brandle, Menna E Jones

**Affiliations:** School of Natural Sciences, University of Tasmania, Private Bag 55, Hobart, TAS 7001, Australia; Vertebrate Pest Research Unit, NSW Department of Primary Industries, Orange, NSW 2800, Australia; Evolution & Ecology Research Centre, School of Biological, Earth and Environmental Sciences, The University of New South Wales, Sydney 2052, Australia; Department for Environment and Water, Port Augusta, SA 5700, Australia; Centre for Ecosystem Science, University of New South Wales, Kensington, NSW 2052, Australia; Davies Livestock Research Centre, School of Animal and Veterinary Sciences, University of Adelaide, Roseworthy, SA 5371, Australia

**Keywords:** invasive species, *Oryctolagus cuniculus*, *Felis catus*, pest management, bottom-up control, conservation, camera survey, BACI, warren ripping, RHDV

## Abstract

The availability of invasive prey often plays an important role in regulating co-introduced invasive predator populations. As predators have been shown to respond rapidly to declines in prey populations, our objective was to experimentally test how local population reduction of an invasive prey species, the European rabbit (*Oryctolagus cuniculus*), affects the activity of an introduced predator, the feral cat (*Felis catus*). To test the effectiveness of three different rabbit control methods, activity levels of cats were surveyed with remote infrared wildlife cameras in three treatment and four control sites. The rabbit control treatments were implemented in extensive open landscapes in the semi-arid zone of South Australia, and consisted of shooting of rabbits, destruction of rabbit warrens, and the targeted delivery of baits treated with RHDV. The results indicate that only the destruction of rabbit warrens has observable effects on the number of cat detections on cameras. Cat detections decreased in the areas where rabbit warrens were destroyed and increased in adjacent areas where rabbits were still abundant. This suggests that cats vacated the treated area and moved into surrounding areas of abundant introduced prey.

## Introduction

Feral cats (*Felis catus*) are a widespread problem in Australia. Their devastating impacts on populations of native animals and the consequences for the functionality of ecosystems are well documented (Woinarski *et al*. 2011; Doherty *et al*. 2016). Efforts to control cats by human intervention have proven to be difficult at large landscape scale outside of fences and islands (Doherty *et al*. 2017). Mostly, cat control is implemented using resource-intensive lethal methods that exert top-down regulation on cat numbers such as shooting, poisoning, and trapping. The resulting overall impact on the population size of feral cats in the target area can be very small (Legge *et al*. 2017). In large open landscapes, such as the Ikara-Flinders Ranges National Park in South Australia (IFRNP), regulating the cat population can be especially difficult, requiring broad-scale methods such as aerial baiting (Moseby *et al*. 2021). Areas where feral cats have been shot, poisoned or trapped are quickly reoccupied (Lazenby, Mooney & Dickman 2014), assisted by their capacity to travel long distances (Jansen *et al*. 2021; Roshier & Carter 2021). It is therefore necessary to implement other or additional solutions to control cats in open landscapes (Denny & Dickman 2010). Ecologically based methods might herby be more effective in reducing cat numbers than lethal control.

Feral cats are opportunistic, generalist carnivores (Bonnaud *et al*. 2010). Their population densities are associated with high prey densities (Pech *et al*. 1992; Holden & Mutze 2002). In Australia, they rely on the invasive European rabbits (*Oryctolagus cuniculus*) as a main food source when rabbits are abundant (Molsher, Newsome & Dickman 1999; Doherty *et al*. 2015). As geographic variation and changes in prey abundance regulate the density of predator populations (Fuller & Sievert 2001), controlling the main food source therefore offers opportunities for bottom-up control of cat abundance (Brothers, Skira & Copson 1985; Mutze 2016; Lurgi, Ritchie & Fordham 2018; McGregor *et al*. 2019). Predators relying on rabbits as a main food source have been shown to respond rapidly to declines in rabbit numbers (Ferreras *et al*. 2011). This was seen after the >90% reduction in rabbits from the initial rabbit haemorrhagic disease (RHDV) outbreak in semi-arid South Australia (Read & Bowen 2001; Holden & Mutze 2002) which showed a response of cats to prey decline and resulted in positive environmental effects for whole ecosystems in arid Australia (Pedler *et al*. 2016).

With the dingo (*Canis dingo*) near-absent, and the red fox (*Vulpes vulpes*) successfully controlled in the IFRNP (Stobo-Wilson *et al*. 2020), any suppressive top-down regulation on feral cats, from intraguild predation or aggressive interference competition is relaxed. Cat abundance is therefore limited by the carrying capacity of their prey (Feit, Feit & Letnic 2019) and the availability of thermal refuge (Briscoe *et al*. 2022). Changing rabbit abundance in an open landscape will create a lower carrying capacity and change the cat density (Pech *et al*. 1992; Fuller & Sievert 2001; Lurgi, Ritchie & Fordham 2018). Recent research simulated this rabbit population decline in an enclosed experimental setup to demonstrate effects on cat activity (McGregor *et al*. 2019). After a short period of prey-switching to sparse other available prey, survival decreased and a large proportion of the feral cats in the treated area starved. In an open landscape, however, changes in prey abundance can have different effects on the behaviour of predators. Feral cats in New Zealand have been observed to alter the size of their home range following a sudden population decline of their primary prey species (Norbury, Norbury & Heyward 1998). The response to a decline in prey abundance is therefore likely to be distinctive in different environments. Ecologists are only starting to understand the consequences of pest management for single species within ecosystems and complex food webs (Ramsey & Norbury 2009).

There is a long history of rabbit control for agricultural purposes in Australia (Berman, Brennan & Elsworth 2011; Cooke *et al*. 2013). Experimental suppression of rabbit populations can be done by a variety of active interventions such as mechanical destruction of warrens by ripping and blasting, broadcasting of poison baits, delivery of RHDV via baits, fumigation of warrens, or shooting (Williams *et al*. 1995). All treatments vary in cost-efficiency and risk to non-target fauna. Until now, most of the research related to rabbit control primarily targets reducing herbivore impact on agriculture, and not on secondary effects like cat control (Smith, Prickett & Cowan 2007; Cooke *et al*. 2013). Even though research has demonstrated the connection between cat and rabbit numbers (Holden & Mutze 2002), knowledge about the efficacy of using bottom-up mechanisms to control cat activity is scarce.

In this paper, our objective is to experimentally test the effectiveness of three different rabbit control methods, implemented in extensive open landscapes, in altering the spatial or temporal activity of feral cats. The treatments were shooting of rabbits, destruction of rabbit warrens, and the spread of baits treated with RHDV. Activity levels of cats are surveyed with remote infrared wildlife cameras in three treatment and four control sites.

## Materials and Methods

### Study Sites

The study was conducted in the Ikara-Flinders Ranges National Park (IFRNP) (31°22’01”S,138°38’34”E) and neighbouring pastoral sheep property Gum Creek Station (31°11’56”S, 138°39’27”E) in seasonally hot, semi-arid South Australia. The study area has low, unreliable rainfall (Morton *et al*. 2011) averaging around 300 mm/year (BOM 2021). European introductions of herbivores to the region have had large impacts on vegetation structure and composition in the Flinders Ranges more broadly, including at these sites (Mutze 2016). We selected seven sites in open grassland (Brandle 2001) that have similar vegetation, habitat structure and geology. Three of the sites were selected as treatment sites and four used as control sites (see Table 1; Figure 1).

**Table 1:** Overview of the survey sites in the Flinders Ranges.

| Survey Site | Size (ha) | Treatment | Year | Days surveyed total | Days surveyed after rabbit control | Eradicat® baited | Fox baited | Previous rabbit control |
| --- | --- | --- | --- | --- | --- | --- | --- | --- |
| Oraparinna South (NP) | 2208 | Shooting of rabbits | 2019 | 167 | 93 | No | Yes | Yes |
| Oraparinna North (NP) | 2249 | Control | 2019 | 167 | - | No | Yes | Yes |
| Gum Creek station West | 1323 | Warren ripping | 2019 | 180 | 90 | No | Yes | Yes |
| Gum Creek station East | 1745 | Control | 2019 | 180 | - | No | Yes | Yes |
| Brachina (NP) | 2143 | RDHV | 2020 | 188 | 143 | Yes | Yes | No |
| Bunyeroo (NP) | 2477 | Control | 2020 | 155 | - | Yes | Yes | No |
| Wilpena (NP) | 2510 | Control | 2020 | 168 | - | Yes | Yes | No |

**Figure 1:**
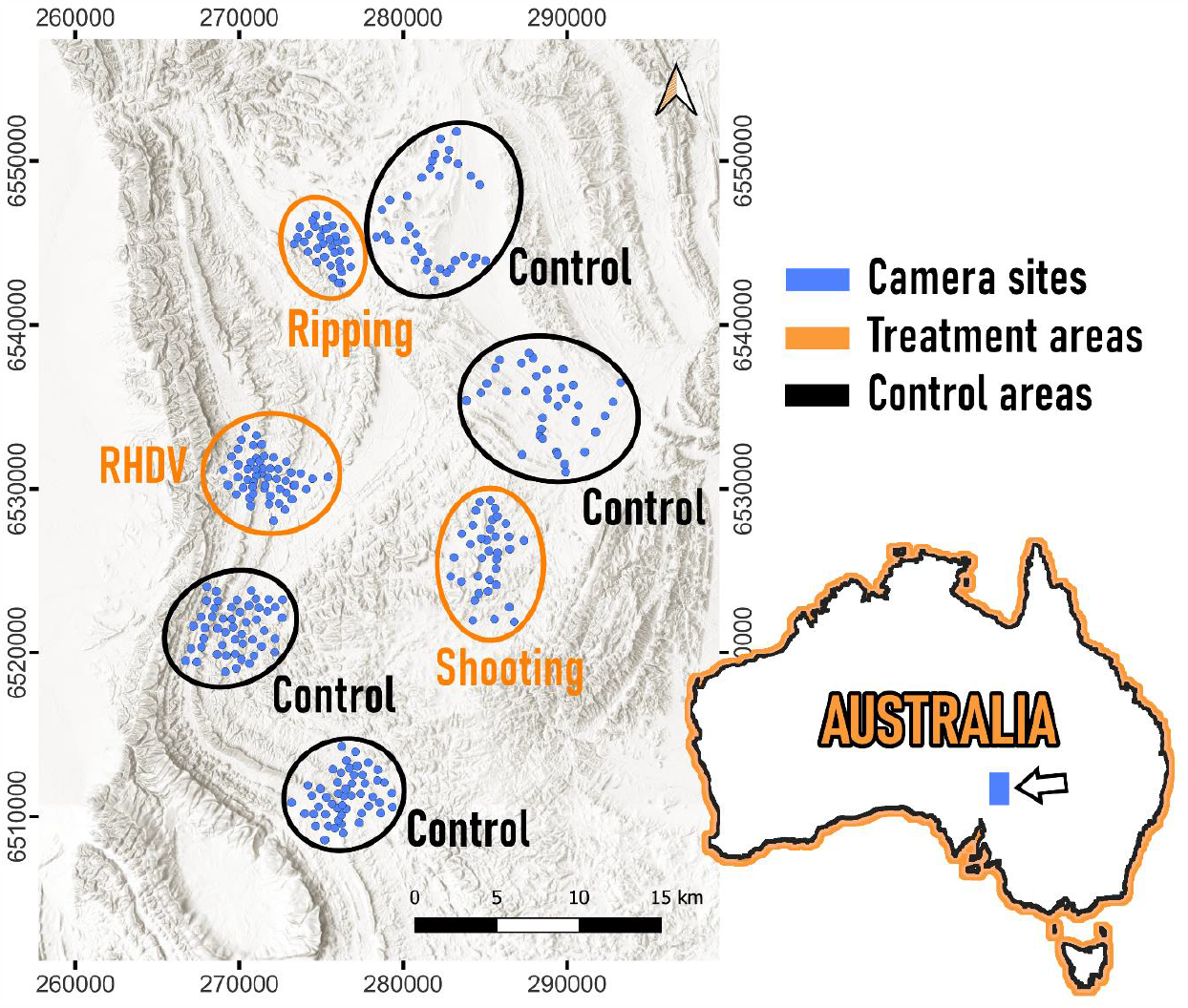
Survey sites in the Flinders Ranges/ South Australia. Blue indicating camera locations, orange the three treatment areas, and in black the control areas. Bottom right inset shows the location within Australia.

Two control and treatment sites were each located in the eastern side of the National Park (Oraparinna North and South) and on Gum Creek station (Gum Creek East and West). These four sites had extensive rabbit control treatments in the late 1990s and have not seen major cat control treatment in recent years (Department of Environment and Heritage 2006). The remaining three sites, one treatment and two control sites, were in the western part of the National Park, near Brachina Gorge, Bunyeroo Gorge and Wilpena Pound entrance respectively. These sites have not had extensive rabbit treatment in the past but have had cat control by spotlight shooting and experimental control with aerial *Eradicat*^®^ baiting.

### Camera surveys

Within each of the four survey areas that had not had major cat control (Gum Creek East and West, Oraparinna South and North), 30 remote infrared cameras (Reconyx PC600 and 800) were set up overviewing creek-lines and paths used by animals. These landscape features had the highest detection rates of cats in past surveys in the I-FRNP and therefore provide the highest accuracy for surveying cat activity in this landscape (Read *et al*. 2015). Additionally, in a central line across each of the four areas, five cameras were set up on active rabbit warrens to record rabbit and cat activity. The three sites with ongoing cat management (Brachina, Bunyeroo, Wilpena) were monitored with 50 cameras set in creeks and minor drainage lines.

Pre-selection of camera sites was done using aerial imagery (Google Earth) and was based on results from previous research in the IFRNP that identified areas with increased cat detection (Stobo-Wilson *et al*. 2020). The mean minimum distance between cameras was 586 m (± 213 m SD). Cameras were set up facing south to avoid direct sun-exposure on the camera lens. Cameras were attached to trees, star-pickets or other solid features at a height of approximately 60 cm and programmed to take a set of five pictures in “balanced” mode with no delay once movement was detected. Functionality of cameras was tested by walking across the area overviewed by the camera at distances of 20 m, 16 m, 12 m, 8 m, 4 m, and 1 m waving a hand at ground level. Only if at least five of the six distances triggered the camera during the test, was the camera declared suitable for the survey. Cameras set on rabbit warrens faced the burrow entrance that had the greatest sign of activity.

Remote cameras were set to collect images of wildlife across the same time periods in both treatment and control sites. Cameras were set for 74 days at the Oraparinna sites and 90 days at the Gum Creek sites before the manipulations occurred. Data collection continued for 93 days at Oraparinna after treatment, and for 58 and 32 days, respectively after first and second treatments at Gum Creek (see Table 1). During this period, the batteries of each camera were changed, and their SD cards downloaded four times. Camera data collection in Brachina, Bunyeroo and Wilpena started in March 2020 and ended at the end of August (188, 155 and 168 days).

### Experimental rabbit treatment

We compared changes in cat detections on camera before and after treatment, comparing three different methods of rabbit control commonly deployed in open landscapes. Data from treatment sites were directly comparable with control sites where cameras were set for the same period. Treatments were ground-shooting of rabbits (Oraparinna South), physical destruction of rabbit warrens by ripping (Gum Creek West), and RHDV baits delivered down burrows (Brachina). The remaining sites were used as control sites (Oraparinna North, Gum Creek East, Bunyeroo and Wilpena).

### Warren ripping

Ripping of rabbit warrens is considered to be the most effective treatment to keep rabbit numbers at low abundance in the long term (Williams *et al*. 1995). Prior to ripping, the rabbit warrens at Gum Creek West were documented as follows: a GPS-point was taken at the centre of the warren, the length and width were measured, and active and inactive entrances were counted. Indicators for activity were fresh rabbit scats, the absence of any obstructions (e.g., spider webs, dried wind-blown plant material) in the burrow entrance, soil-scratches, and signs of digging. All warrens in the area were recorded even though some could not be ripped (e.g., slope too steep). We recorded whether a warren had previously been ripped, independent of the number of entrances. Warren ripping at Gum creek was undertaken from 22-28/7/2019 and again from 24-27/9/2019 as part of pest management undertaken by the pastoralist. Starting in the east of the survey area and continuing clockwise, 187 warrens were destroyed during the first ripping period, and a further 85 in the second ripping period. Ripping was done using a loader with a ripper comprising three 0.9 m long tines mounted to the front. The loader was driven backwards, starting 0.5-1 m further out beyond all entrances of the warren. After an initial parallel ripping across the whole warren, the warren was then cross-ripped at approximately 90 degrees to the initial rip-lines (Williams *et al*. 1995).

### Shooting of rabbits

Shooting of rabbits can very rapidly reduce rabbits to low numbers but is considered an ineffective method to control rabbit populations in the long-term (Williams *et al*. 1995). Shooting also allows keeping track of the number of rabbits removed. Rabbit control by shooting at Oraparinna South was undertaken from 4-10/7/2019 and again from 21-28/7/2019 by a professional contractor. Rabbits were shot at night from an all-terrain vehicle equipped with a spotlight, with carcasses left in situ. Sightings of rabbits, and the number of rabbits that were shot were recorded with a GPS data-logger (Keskull/ Evo 2 Muti Event). The entire area was covered regularly by the contractor, but the focus of the shooting effort was on areas with higher numbers of rabbit sightings.

### RHDV treatment

The original serotype of Rabbit Haemorrhagic Disease Virus (RHDV1) was established accidentally in Australia in October 1995 and in most areas resulted in a dramatic decline in the abundance of rabbits (Mutze, Cooke & Alexander 1998). However, the effectiveness of this virus decreased over time and a new Korean variant, RHDVa-K5, was released by state government authorities and landholders in 2017 (Cox *et al*. 2019). This serotype was used as the RHDV treatment in IFRNP, with introduction of the virus and initiation of an outbreak only requiring ingestion of a few viral particles by some susceptible rabbits. Prior to its experimental release by the Department of Environment and Water (DEW) during this study, two sets of prefeeding with 400 kg of carrot pieces took place on 9-10/4/2020 and 14-15/4/2020. Carrots were chopped up into pieces approximately 3 cm x 3 cm x 3 cm. Free feeding comprised laying a line of chopped carrots across all main mapped warren complexes. To reduce non-target uptake by kangaroos and improve rabbit exposure to the carrots, the subsequent free-feed and virus loaded bait placed most of the carrot into the most active burrow entrances.

To prepare the virus loaded bait, 10 ml of reconstituted RHDVa-K5 virus was diluted with 90 mL of water and mixed in a cement mixer with 10 kg of chopped carrots. Each 10 kg of this mix was then bagged into plastic bags. Treated carrots were distributed on 19-20/4/2020 into 76 of the biggest rabbit warrens in the Brachina area.

### Analysis of camera images

Camera images were analysed using *Reconyx MapView Professional* 3.7.2.2. Images of each individual animal were tagged according to their species using the clearest image from the five-image burst. Each animal was tagged only once. If more than one animal was present in an image, successive images were tagged to account for all the animals. To avoid double counting, individual animals were only tagged after a separation time of 10 minute between captures consistent with other studies in the area (e.g. Dean *et al*. 2023).

### Analysis of camera image data

All analyses were done in *R* using *RStudio* 2021.09.0 (RStudio Team 2020). The information for each individual camera included the area, treatment type, and the week of year to account for seasonality. The number of detections of cats and rabbits per camera was then pooled for each week. Many nocturnal animals are known to adapt their activity according to lunar illumination (Daly *et al*. 1992; Pratas-Santiago *et al*. 2017). We therefore included the average illuminated fraction of the moon and its phase per week calculated from the daily values using the package *suncalc* (Thieurmel, Elmarhraoui & Thieurmel 2019).

The detections of cats on the individual cameras were analysed using a general linear mixed model with the package *glmmTMB* (Magnusson *et al*. 2017). Individual camera ID and area were included as a random factor. Model selection was done using Akaike’ s information criterion, with ΔAIC ≤ 6 used to select the most parsimonious model, as in Richards (2008).

### Analysis of rabbit densities for the maps

Rabbit densities were calculated using warren locations and the heatmap plugin in *QGIS* (QGIS Development Team 2020) using the total number of holes as the weight and 200 m as the distance of spread from the centre of the warren. This value is based on the home range size of rabbits and their impact on the vegetation surrounding their warrens (Lange & Graham 1983; Moseby *et al*. 2005). In Gum Creek West we calculated rabbit densities using the data from the mapped warrens, progressively removing warrens from the density estimates as they were ripped during the treatment. In Oraparinna South we used the location where the rabbits were shot to indicate the spatial spread of the management effort.

## Results

### *Warren* ripping

Of the whole treatment site at Gum Creek West (1323 ha), 524 ha or 39.6% of the area was treated during the survey period with ripping in two sessions. During the first ripping period (4 days), 187 warrens with 2140 active and 1750 inactive holes were destroyed. A further 85 warrens were destroyed in the second ripping period (2 days), comprising 1022 active and 849 inactive holes. Together these comprised 71.6% of the active holes and 61.7% of the inactive holes at this site. The detections of rabbits started to decrease at the end of July in Gum Creek West while cat detections steadily rose from the end of August (Figure 2; Gum Creek West).

**Figure 2:**
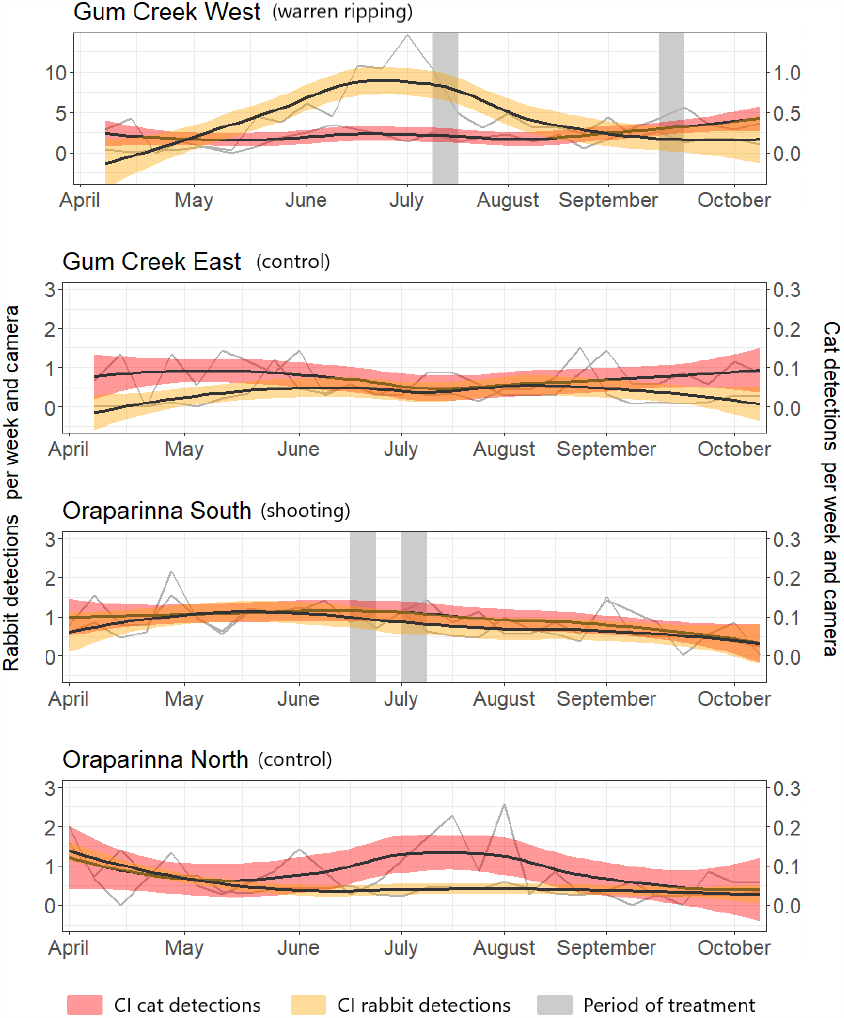
Cat (red) and rabbit (yellow) detections on camera at Gum Creek West (warren ripping), Oraparinna South (shooting of rabbits), Gum Creek East and Oraparinna North (both control areas). Confidence intervals (CI) are displayed in colour. The grey bars indicate the time of treatment.

### Shooting of rabbits

During the treatment period, the contractor spent 12 nights shooting rabbits with a total of 855 rabbits detected of which 703 were shot. Rabbit and cat detections stayed relatively stable throughout the study period in the area of the shooting (Figure 2; Oraparinna South). A minimal decline of rabbit detections is observed after the end of June with cat detections slightly decreasing at the end of July.

### RHDV treatment

Approximately 400kg RHDV treated carrots were distributed on 19-20/4/2020 into 76 rabbit warren entrances in the Brachina area. After the treatment at Brachina, rabbit detections increased until the end of July and decreased again thereafter (Figure 3; Brachina). Cat detections in the area were stable with a slight peak at the end of June.

**Figure 3:**
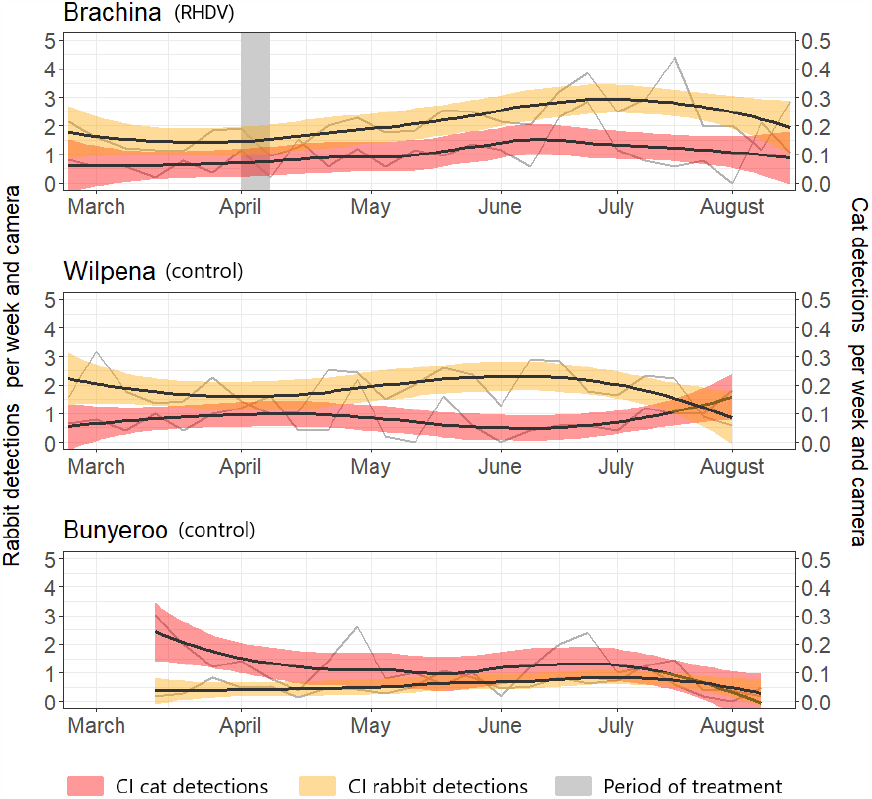
Cat (red) and rabbit (yellow) detections on camera at Brachina (RHDV treatment), Bunyeroo and Wilpena (both control areas). Confidence intervals are displayed in colour. The grey bars indicate the time of treatment.

### Control sites

Both Gum Creek East and Oraparinna North had stable rabbit detections throughout the study period (Figure 2). Cat detections at Gum Creek East stayed level, whereas Oraparinna North had a slight rise in cat detections between June and September. Rabbit detections both in Wilpena and Bunyeroo were stable through the study until July after which they decreased (Figure 3). Cat detections in these areas were also stable with a slight increase in the Wilpena and a slight increase in the Bunyeroo area after July.

### Relative effectiveness of the treatments

The final model describing cat activity included the terms: number of rabbit detections, moon fraction and the interaction between treatment type and management (BACI). The results of the most parsimonious model show that cats were more likely to be detected on cameras that have a higher number of rabbit detections (Figure 4a and c). There was a very weak trend of higher detections of cats with higher illuminated fraction of the moon (Figure 4b and c). Ripping of rabbit warrens had a strong influence on cat detections, there was an effect of the spread of RHDV, and shootings was not influencing cat detections (Figure 4c and d). Week of the year and moon-phase (waxing/ waning) did not explain cat detections and were removed from the final model.

**Figure 4:**
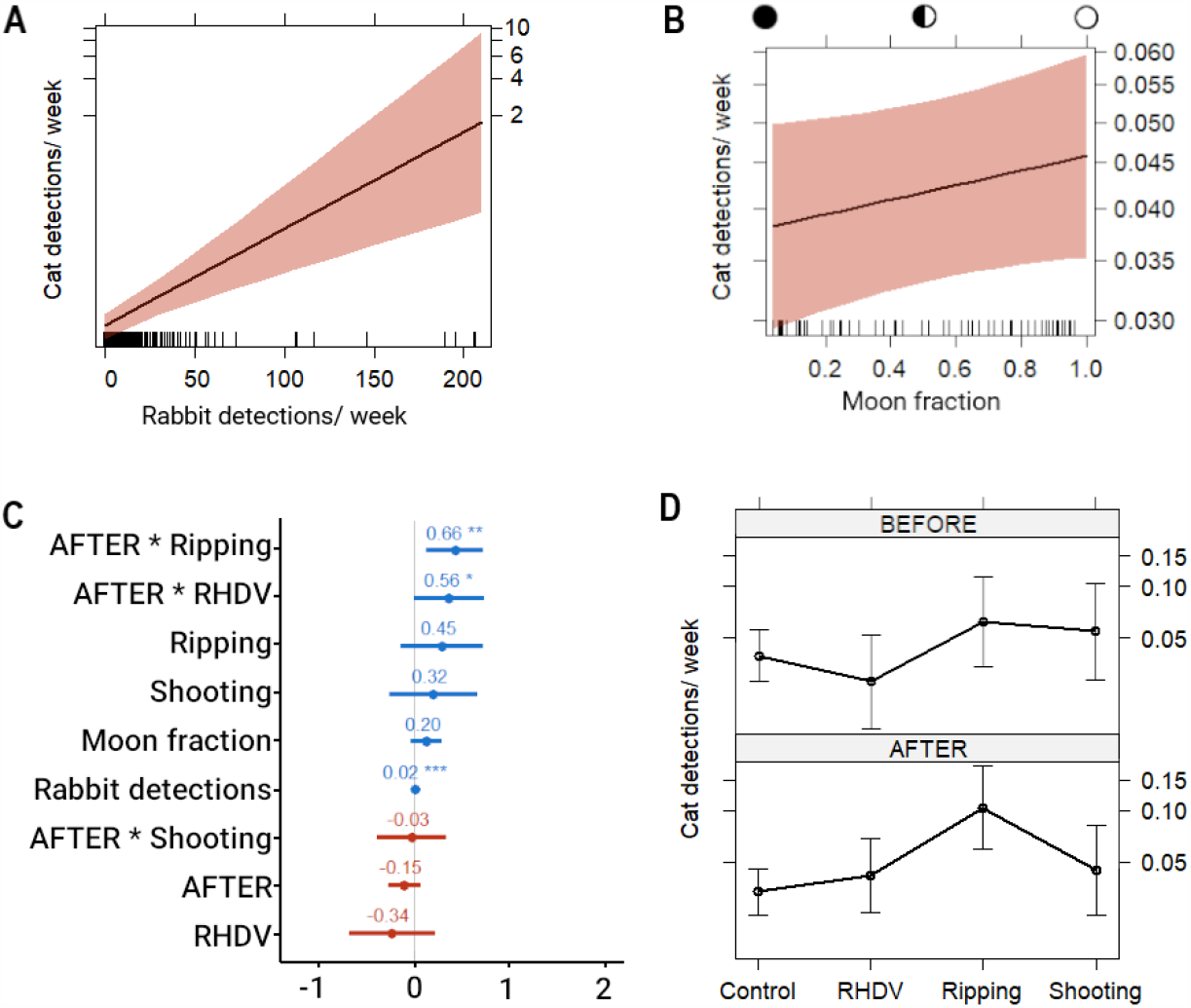
Effects of the best model describing the influence of rabbit control treatments on rabbit and cat detections on camera: a) detection of cats in relation to the detection of rabbits across all camera sites, b) effect of the illuminated fraction of the moon on cat detection, c) estimates and standard error of the final model d) effects of different treatments on the number of cat detections per week. Confidence intervals in (a) and (b) are displayed in colour.

The spatial distributions of cat detections within the site treated by ripping shifted following the treatment. Following the progressive warren destruction across the site, cats were detected more frequently overall, but detections reduced to zero in the treated parts and were much higher in the adjacent untreated parts of the site which still had high rabbit densities (Figure 5). In the shooting treatment area, a shift in the spatial detections of cats was not evident. Numbers of detections remained at a similar level, although a minor redistribution of detections from a high number of detections on a few cameras to fewer detections on a greater number of cameras can be observed.

**Figure 5:**
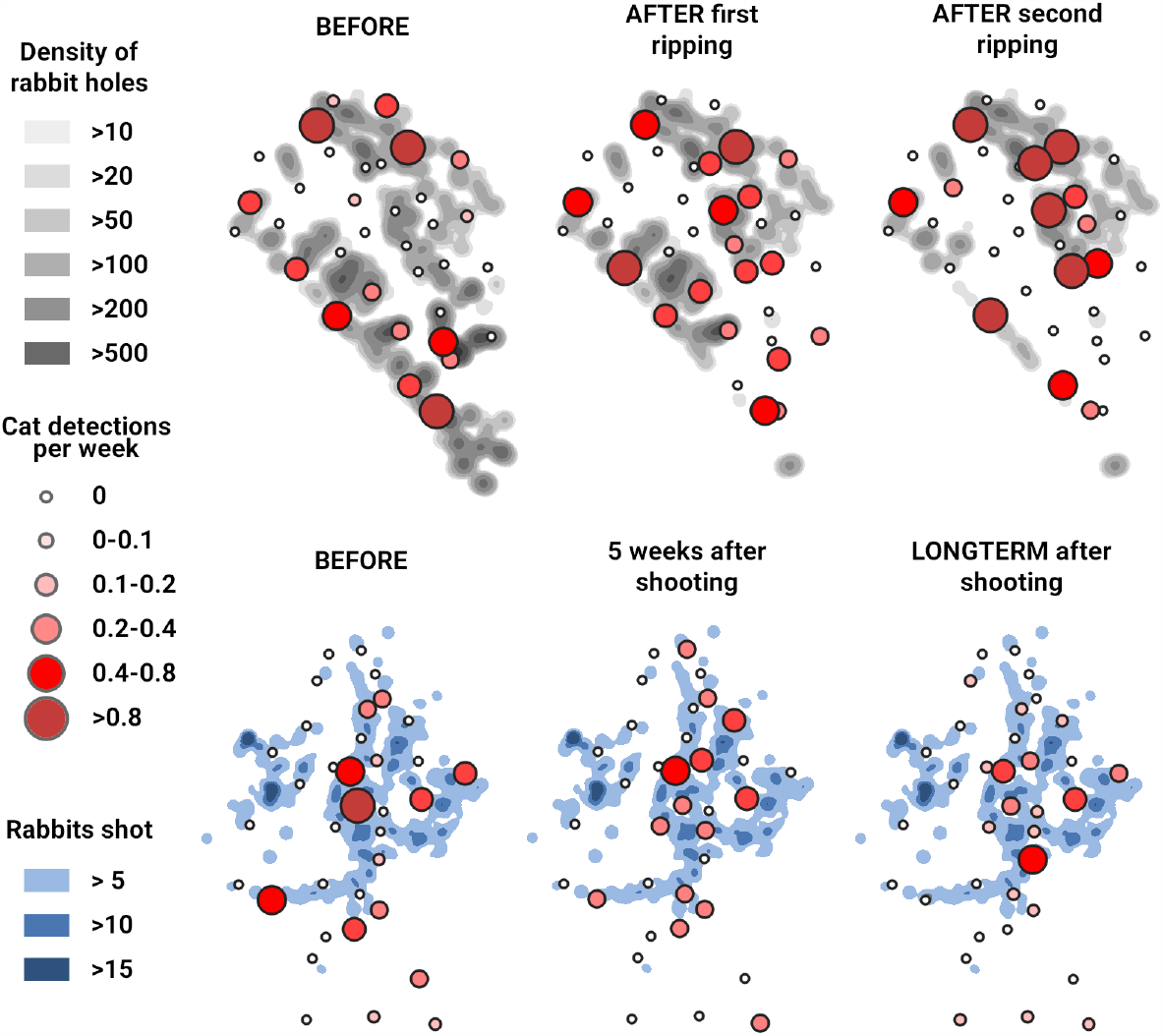
Effect of rabbit removal by ripping (top row) and shooting (bottom row) on weekly detection of cats on camera (in red). Polygons displayed are rabbit hole densities (in grey) and rabbits shot during pest management (in blue).

## Discussion

Our study shows that the spatial activity of feral cats can be influenced by reducing the local abundance of rabbits, where rabbits are their primary prey, and that the mechanical destruction of rabbit warrens is more effective in influencing the magnitude of the outcome than shooting or baiting with RHDV as a method of rabbit control. The number of cat detections in the study areas is strongly connected to the number of rabbit detections. When the local numbers of rabbits were severely reduced, cat activity shifted to adjacent areas where rabbit density was still high. Any reduction of rabbit numbers should therefore have an effect on cat activity. Our study shows that warren ripping, identified in previous research as the most efficient way to reduce rabbit numbers (McPhee & Butler 2010), also had the greatest effect on cat activity. Effective rabbit treatment consequently can be used by conservation managers to influence cat presence. This could be exploited to connect areas of suitable habitat if cats can be moved out of strategic areas to reduce threats from predation and allow occupation or movement of native animals.

While the warren ripping resulted in lower rabbit detections, results suggest neither the shooting nor the RHDV release were successful in affecting rabbit or cat detections on camera. This is despite a high percentage of observed rabbits being removed by the shooter. The threshold of numbers of rabbit removed seems to be high before effects on cat activity are detectable using camera survey data. This may be due to generally low detection rates of cats where changes are less obvious or indicate a high number of unobserved rabbits, likely within burrows. Results from the release of RHDV down burrows are similar to those of shooting, with insufficient rabbits removed to make a difference in cat detections. The spread of RHDV relies on the interaction between rabbit susceptibility to the virus and insect vectors (Mutze *et al*. 2002; Mutze *et al*. 2014). An effective spread and suppression of rabbits by artificially introducing RHDV is therefore difficult to achieve (Mutze *et al*. 2010). The dry conditions during the study period, resulting in lower fly abundance, might have precluded the ideal conditions required for the spread of RHDV (Asgari *et al*. 1998). Another factor mitigating against the effectiveness of RHDV in reducing rabbit densities in the study is the previous outbreaks of RHDV1 and RHDV2 in this landscape. These outbreaks may have provided the rabbits with some evolutionary resistance against the RHDVa-K5 strain used in this study, thus reducing its impact (Schwensow *et al*. 2017).

Declines in rabbit detections resulted in a rise of cat activity across the study area as a whole during our study. This rise in cat detections, especially in the site where we ripped warrens, is counterintuitive. A detailed examination of the spatial component of cat detections, however, indicate that cats moved out of the treated areas, sequentially as areas of warrens were ripped. This effect is clearly visible in the ripping site of Gum Creek West. It is less evident in the site where rabbit shooting occurred, where elevated cat detections appear to be spread out over a larger area. Reducing their main food source appears to have forced the cats to higher mobility in the search for prey, resulting in the increased detections seen in adjacent areas that still had high rabbit densities.

Our result indicate that feral cats prefer to move into nearby areas of preferred habitat if pushed out of their original surroundings. We found no evidence of cats remaining in the ripped area. If they had stayed in the ripped area this could have been interpreted as a sign that they were able to switch to alternative prey species. Such a dietary change was seen at Astrebla Downs National Park (Queensland) where feral cats switched to bilbies (*Macrotis lagotis*) when numbers of the hyperabundant longhaired rats (*Rattus villosissimus*) dramatically reduced at the end of a population irruption (Rich *et al*. 2014). There was no indication in our study of behaviours consistent with prey switching, noting a comparable prey species to rabbits is not present where the ripping was done. Our results support the research indicating that in the absence of alternative prey cats will move, if possible, or die (Read & Bowen 2001; McGregor *et al*. 2019).

## Conclusions

We show that, in arid zone environments, reducing local rabbit populations can shift the activity of cats out of areas of former high rabbit abundance. This presents an opportunity for conservation managers to apply rabbit control as an approach to influence feral cat presence, particularly when rabbits serve as the primary prey. Our results indicate that a high threshold of rabbit removal is needed to achieve cat activity changes. Mechanical warren destruction, previously established as highly effective in reducing rabbit numbers, has the most pronounced impact on cat activity, while RHDV release had a minor effect and rabbit shooting did not significantly affect rabbit or cat detections. Declining rabbit numbers led to increased cat activity across the study area, with cats relocating from treated areas, particularly in the warren-ripped site. In essence, our study underlines that feral cats move to nearby areas with high rabbit abundance habitats rather than staying in treated zones. These findings support the idea that feral cats will move to secure their main food source, reinforcing the importance of effective rabbit control in shaping feral cat behaviour and their spatial distribution. This might facilitate connecting suitable habitats and therefore mitigate predation risks to native wildlife.

## Author Contributions

JJ, KEM, DEP, RB and MEJ conceived the ideas and designed methodology; JJ, ATD, GA and RB collected the data; JJ, SC and ATD analysed the data; JJ, DEP, RB and MEJ led the writing of the manuscript. All authors contributed critically to the drafts and gave final approval for publication.

## Acknowledgements

We thank Bill McIntosh and family for their support, permission to conduct fieldwork on Gum Creek Station, and undertaking the warren ripping; Greg Mutze for suggestions on study design; the Department for Environment and Water for support during the study; Adnyamathanha members of the IFRNP co-management board for permission to do fieldwork on their country; National Parks and Wildlife Service for ongoing support and access to park facilities; Frank Bernhardt (Bernhardt Pest Control) for undertaking the rabbit shooting; Cat Lynch for support in the early stages of the project; Leon Barmuta and people in the School of Natural Sciences for help during analysis and Jan Jansen for consistent support.

The study was funded by an Australian Research Council Discovery grant (DP170101653 to MJ). Jeroen Jansen was supported by a Tasmania Graduate Research and a Tuition Fee Scholarship.

## Data Availability Statement

The data presented in this study will be available on request from the Department for Environment and Water South Australia

